# Apparent RNA bridging between PRC2 and chromatin is an artefact of non-specific chromatin precipitation upon RNA degradation

**DOI:** 10.1101/2023.08.16.553503

**Authors:** Alex Hall Hickman, Richard G. Jenner

## Abstract

Polycomb repressive complex 2 (PRC2) modifies chromatin to maintain repression of genes specific for other cell lineages and is essential for development. In vitro, RNA inhibits PRC2 catalytic activity and interaction with nucleosomes but the effect of RNA on PRC2 in cells has been unclear, with studies concluding that RNA either antagonises or promotes the association of PRC2 with chromatin. The addition of RNase A to chromatin immunoprecipitation reactions has been reported to reduce detection of PRC2 target sites, suggesting the existence of an RNA bridge that connects PRC2 to chromatin. Here, we show that this apparent loss of PRC2 chromatin association after RNase A treatment is due to non-specific chromatin precipitation. RNA degradation causes chromatin to precipitate out of solution, thereby reducing the enrichment of specific DNA sequences in chromatin immunoprecipitation reactions. Maintaining chromatin solubility by the addition of poly-L-glutamic acid (PGA) rescues detection of PRC2 chromatin association upon RNA degradation. These findings undermine support for the model that RNA bridges PRC2 and chromatin in cells.

## INTRODUCTION

Chromatin regulation is fundamental for cell identity and cell differentiation. Chromatin modifiers open chromatin at genes important for the cell’s identity and close chromatin at genes specific for other cell types and this is regulated dynamically during cell differentiation. Dysregulation of these processes cause developmental disorders and cancer ^1,2^. Chromatin is associated with a host of RNA species, including nascent RNAs in the process of transcription and a subset of mature transcripts that remain localised to the nucleus ^3–5^. It has become apparent that RNA interacts with chromatin modifiers and modulates their association with chromatin but the mechanisms underlying these effects remain poorly understood ^4–9^

Polycomb repressive complex 2 (PRC2) has become a paradigm for RNA binding chromatin regulators ^4–9^. PRC2 associates with genes specific for other cell lineages and maintains them in a silent state ^10,11^. PRC2 is recruited to chromatin through the interaction of accessory subunits with ubiquitinated H2A (H2AK119ub) and with GC-rich sequences located at CpG islands ^10,11^. The PRC2 enzymatic subunit, EZH2, trimethylates histone H3 at lysine 27 (H3K27me3), which creates a binding site for canonical forms of PRC1 and allosterically activates PRC2 activity through its subunit EED.

That PRC2 also binds RNA was first reported for certain long non-coding RNA (lncRNA) species, which were proposed to recruit PRC2 to specific sites on chromatin ^12–15^. Subsequently, PRC2 was found to interact primarily with nascent pre-mRNAs in cells with a preference for G-tract sequences, especially when folded into G-quadruplex structures ^16–22^. In vitro, RNA inhibits PRC2 catalytic activity ^22–24^ by antagonising its interaction with DNA ^25^, nucleosomes ^17^ and the substrate H3 tail ^19^, and by blocking allosteric activation through EED ^26^.

The effect of RNA on PRC2 function in cells has been less clear. Consistent with in vitro data, tethering G-tract RNA to polycomb target genes reduces PRC2 occupancy and depletes H3K27me3 in cis ^19^. Other studies have concluded that RNA can also promote PRC2’s repressive effects on chromatin in cells ^27,20^. One of the seemingly strongest pieces of evidence that RNA plays a positive role in PRC2 chromatin association in cells are from experiments by Long and colleagues utilising a variant of chromatin immunoprecipitation coupled with sequencing (ChIP-seq) methodology, termed rChIP ^27^. The authors found that the addition of RNase A during the IP step abrogated detection of PRC2 chromatin occupancy but had no effect on RNA polymerase II or TBP. These data were interpreted as RNase A severing an RNA ‘bridge’ that tethered PRC2 to its target sites on chromatin.

To understand further the mechanism by which RNA regulates PRC2 chromatin occupancy, we sought to replicate the rChIP experiments performed by Long et al. We found that the apparent loss of PRC2 chromatin occupancy that occurs upon RNase A treatment stems from non-specific precipitation of chromatin out of solution rather than the loss of PRC2-bound genomic fragments. Our results thus undermine support for the model that RNA bridges PRC2 and chromatin in cells.

## RESULTS

### The RNase A treatment step in rChIP methodology precipitates chromatin from solution

In seeking to understand the basis for the results of Long et al, we performed rChIP for histone H3 in mouse embryonic stem cells (mESC). Sonicated crosslinked cell extracts were prepared as described ^27^ (Long, Rinn and Cech personal communication), and anti-histone H3 antibody added, before the sample was split in two, RNase A added to one of the tubes and both samples incubated overnight. After the subsequent addition of protein A/G beads to the tubes we noticed a clear aggregation of the beads in the sample treated with RNase A (Fig. 1a). Investigating this phenomenon further in the absence of antibody and beads, we found that RNase A treatment caused precipitation of material from the extracts that could be visualised as a large pellet after centrifugation (Fig. 1b). SDS-PAGE demonstrated that this insoluble, pelleted material was enriched in histones, which were correspondingly depleted from the supernatant by RNase A treatment (Fig. 1c). These results indicate that RNase A treatment of sonicated cell extracts in the rChIP procedure causes the precipitation of chromatin from solution.

**Figure 1.**
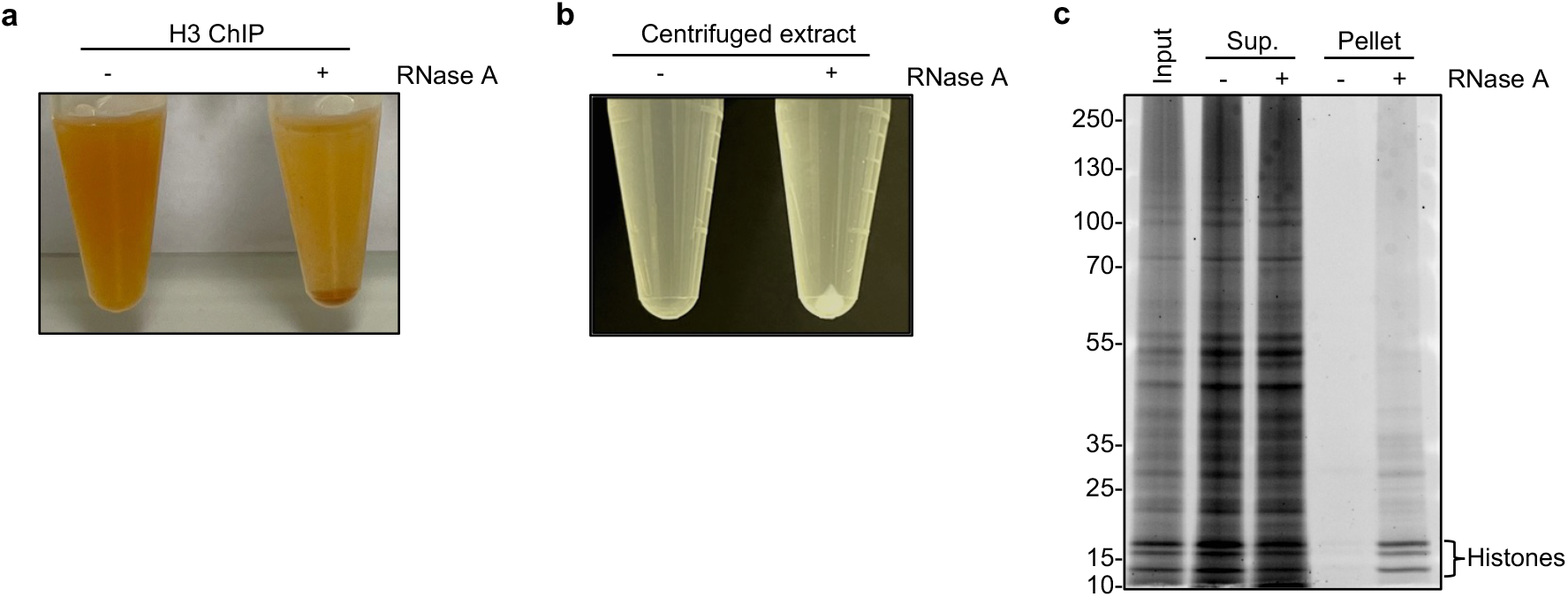
The RNase A treatment step in rChIP methodology precipitates chromatin from solution. **a**. Aggregation and sedimentation of protein A/G magnetic beads during rChIP for histone H3 in RNase A treated sonicated crosslinked mESC extract. Representative of 2 independent experiments. **b**. Presence of an insoluble pellet after centrifugation of sonicated mESC extract prepared according to rChIP methodology following incubation with RNase A. Representative of 2 independent experiments. **c**. SDS-PAGE of equal volumes of input, supernatant (sup.) and pelleted fractions isolated from centrifuged sonicated crosslinked mESC extract treated or not treated with RNase A. Histones are marked. Representative of 2 independent experiments.

### Poly-L-glutamic acid maintains chromatin solubility upon RNA degradation

RNA degradation caused by RNase A treatment has previously been reported to cause precipitation of histones from cell extracts due to electrostatic effects ^28,29^. It was also found that histone solubility could be restored in these conditions by the addition of poly-L-glutamic acid (PGA) ^28^, which like RNA is an anionic polymer. We therefore tested whether the addition of PGA would prevent chromatin from precipitating during the rChIP protocol. We found that addition of PGA immediately prior to RNase A prevented precipitation of proteins, including histones upon RNase A treatment (Fig. 2a). We also found that RNase A treatment precipitated DNA from solution and that this was prevented by PGA addition (Fig. 2b). PGA had no effect on the ability of RNase A to degrade RNA in nuclear extracts (Fig. 2c), demonstrating that PGA did not maintain chromatin solubility by inhibiting RNase A activity.

**Figure 2.**
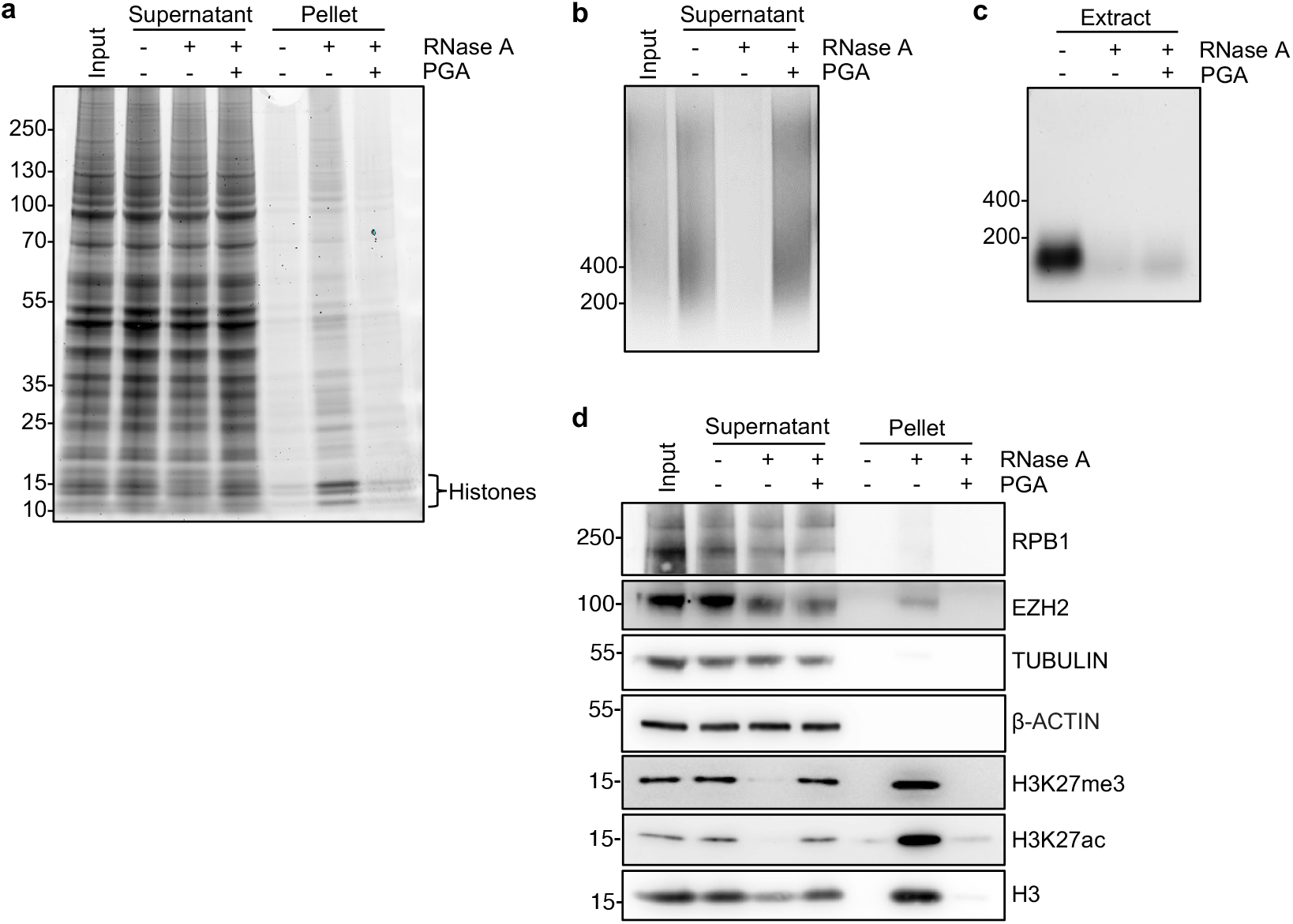
PGA maintains chromatin solubility upon RNA degradation. **a**. SDS-PAGE of equal volumes of input, supernatant and pelleted fractions isolated from centrifuged sonicated mESC extract treated with RNase A alone, RNase A and PGA (400 ng/μl), or untreated. Representative of 2 independent experiments. **b**. Agarose gel electrophoresis of equal volumes of DNA purified from the supernatants of the samples described in a. Representative of 2 independent experiments. **c**. Agarose gel electrophoresis of RNA purified from sonicated mESC extract treated with RNase A alone, RNase A and PGA (400 ng/μl), or untreated. The dashes mark the position of dsDNA size markers. **d**. Immunoblots for the indicated proteins in the samples described in a. Representative of 2 independent experiments.

To characterise the nature of the RNase A induced insoluble material further, we subjected the pellet and supernatant to immunoblotting (Fig. 2d). This confirmed that the insoluble material was enriched for histone H3, including H3 trimethylated at lysine 27 (H3K27me3), and contained the PRC2 subunit EZH2. The RNA polymerase II subunit RBP1 was also present in the pellet but was not as enriched as EZH2 or H3. Furthermore, PGA greatly reduced the precipitation of histone H3, EZH2 and RBP1 caused by RNase A treatment. These results demonstrate that the RNA degradation step in the rChIP protocol cause chromatin and its associated proteins to precipitate from solution and that the addition of an alternative anionic polymer in the form of PGA prevents this from occurring.

### RNase A treatment reduces the specificity of IP reactions

We next tested whether chromatin precipitation in the presence of RNase A affected IP specificity. We performed co-immunoprecipitation (co-IP) for histone H3, SUZ12 and RBP1 in the presence and absence of RNase A and blotted for these factors in the input and IP fractions (Fig. 3a). In the absence of RNase A, each IP reaction specifically enriched its target protein, as expected. However, in the presence of RNase A, the IP reactions exhibited reduced specificity, pulling down other factors. In the presence of RNase A, IP for H3 enriched for EZH2, H3K27me3 and H3K27ac, while IP for the PRC2 subunit SUZ12 and RBP1 also enriched for H3, H3K27me3 and H3K27ac (Fig. 3a). The addition of PGA with RNase A maintained the specificity of each IP. Thus, these data show that the precipitation of chromatin out of solution caused by RNase A in the rChIP procedure reduces the specificity of IP reactions by increasing the amount of background material that is co-precipitated.

**Figure 3.**
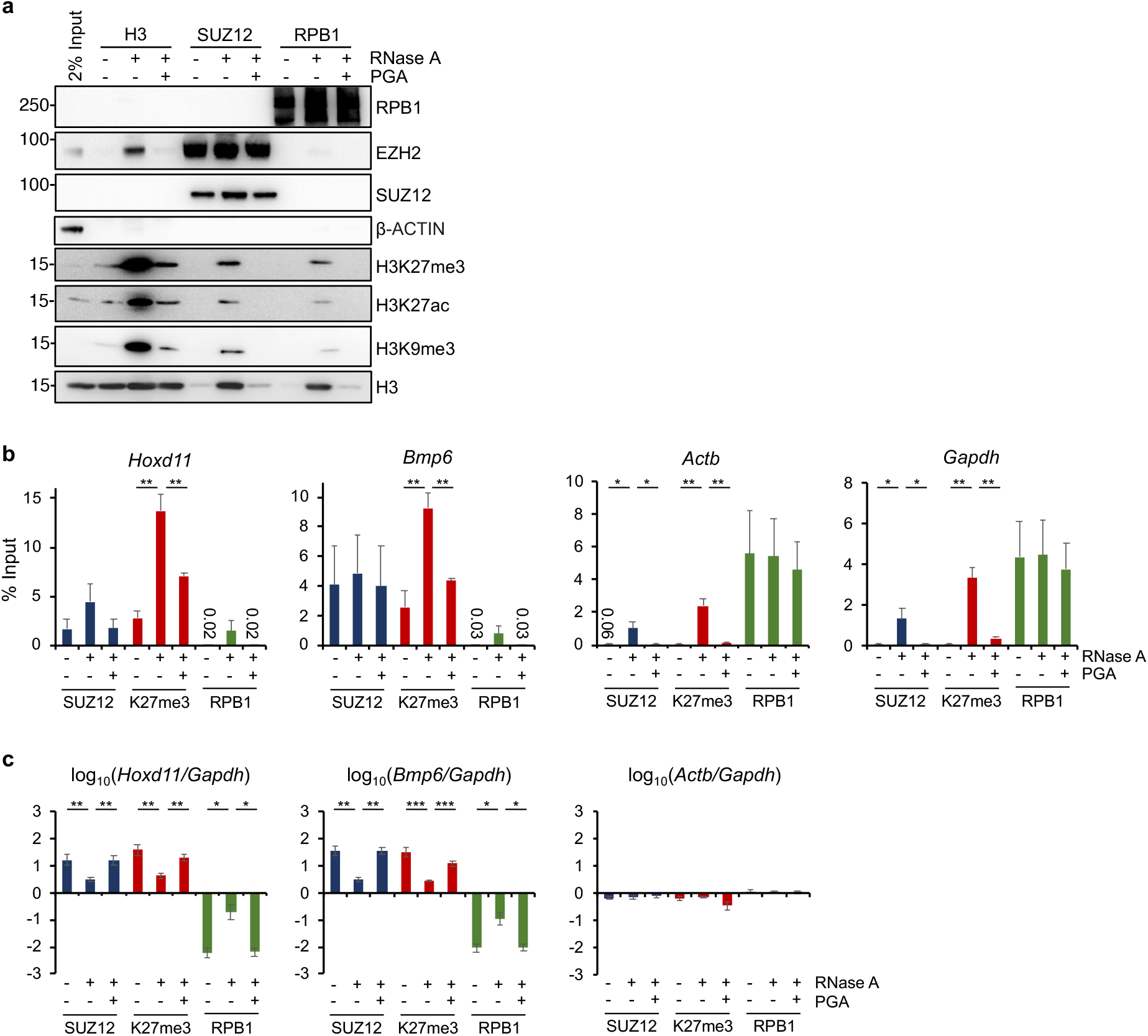
RNase A treatment reduces the specificity of IP reactions. **a**. Immunoblots of the indicated proteins in input and H3, SUZ12 and RPB1 IPs from sonicated crosslinked mESC extracts after treatment with RNase A, RNase A and PGA or left untreated. Representative of 2 independent experiments. **b**. Quantitative PCR (as % input) for *Hoxd11, Bmp6, Actb* and *Gapdh* in SUZ12, H3K27me3 and RBP1 rChIP samples from mock, Rnase A, or RNase A and PGA treated sonicated crosslinked mESC extracts (mean and S.E., 3 independent experiments; *p<0.05, **p<0.01, ***p<0.001, one-sided t-test). **c**. Data in b. except normalised to *Gapdh*.

We next assessed how RNase A treatment impacted the specificity of enrichment of DNA by ChIP. We performed rChIP for SUZ12, H3K27me3 and RBP1 and measured enrichment of the repressed PRC2 target genes *Hoxd11* and *Bmp6* and the active housekeeping genes *Actb and Gapdh* by qPCR (Fig. 3b). In the absence of RNase A, we found that ChIP for SUZ12 and H3K27me3 enriched for *Hoxd11* and *Bmp6*, but not for *Actb or Gapdh*, as expected. Reciprocally, ChIP for RBP1 enriched for *Actb* and *Gapdh* but not for *Hoxd11* and *Bmp6*. Then, examining the enrichment of these genes in samples treated with RNase A, we found increased precipitation of *Hoxd11* and *Bmp6* in H3K27me3 ChIPs with a smaller effect for SUZ12. However, this was coupled with a large increase in precipitation of *Gapdh* and *Actb* in both SUZ12 and H3K27me3 ChIPs. Reciprocally, RNase A treatment increased precipitation of *Hoxd11* and *Bmp6* by RBP1 ChIP, although this did not reach significance. Calculating the ratio of *Hoxd11* and *Bmp6* versus *Gapdh* as a measure of ChIP specificity demonstrated that RNase A treatment significantly reduced the specificity of SUZ12, H3K27me3, and RBP1 ChIPs (Fig. 3c). RNase A treatment reduced the fold enrichment of *Hoxd11* and *Bmp6* versus *Gapdh* in SUZ12 and H3K27me3 ChIPs from between ∼20-60-fold to ∼3-5-fold. Reciprocally, RNase A treatment increased enrichment of *Hoxd11* and *Bmp6* versus *Gapdh* in RBP1 ChIPs. In the presence of PGA, the specificity of each ChIP was substantially restored. We conclude, consistent with our co-IP data, that RNase A treatment in the rChIP protocol reduces the specify of IP reactions by increasing the background, non-specific precipitation of chromatin.

### Apparent RNA-dependent PRC2 occupancy in rChIP experiments is due to non-specific chromatin IP

We considered that the non-specific precipitation of chromatin caused by RNase A treatment could account for the apparent RNA-dependent PRC2 occupancy observed by Long et al. using rChIP ^27^. We hypothesised that if this was the case then RNase A treatment should have the same apparent effect on H3K27me3 occupancy, even though this histone modification is not considered to be attached to chromatin via an RNA bridge. We also hypothesised that the apparent effects of RNase A on PRC2 and H3K27me3 chromatin occupancy should be largely prevented by the addition of PGA.

To test these theories, we performed rChIP for SUZ12 and H3K27me3, and also for RBP1, used by Long et al. as a presumed RNA-independent control. We found that RNase A treatment resulted in the loss of enrichment of PRC2-occupied sites (Figs. 4a, 4b and S1), mimicking the apparent RNA-dependent SUZ12 and EZH2 occupancy reported by Long et al. We also found that RNase A treatment had an identical effect on H3K27me3, almost completely abrogating detection of H3K27me3-occupied sites (Figs. 4a, 4b and S1). Strikingly, the addition of PGA prevented the apparent loss of SUZ12 and H3K27me3 chromatin occupancy caused by RNase A treatment (Figs. 4a, 4b and S1).

**Figure 4.**
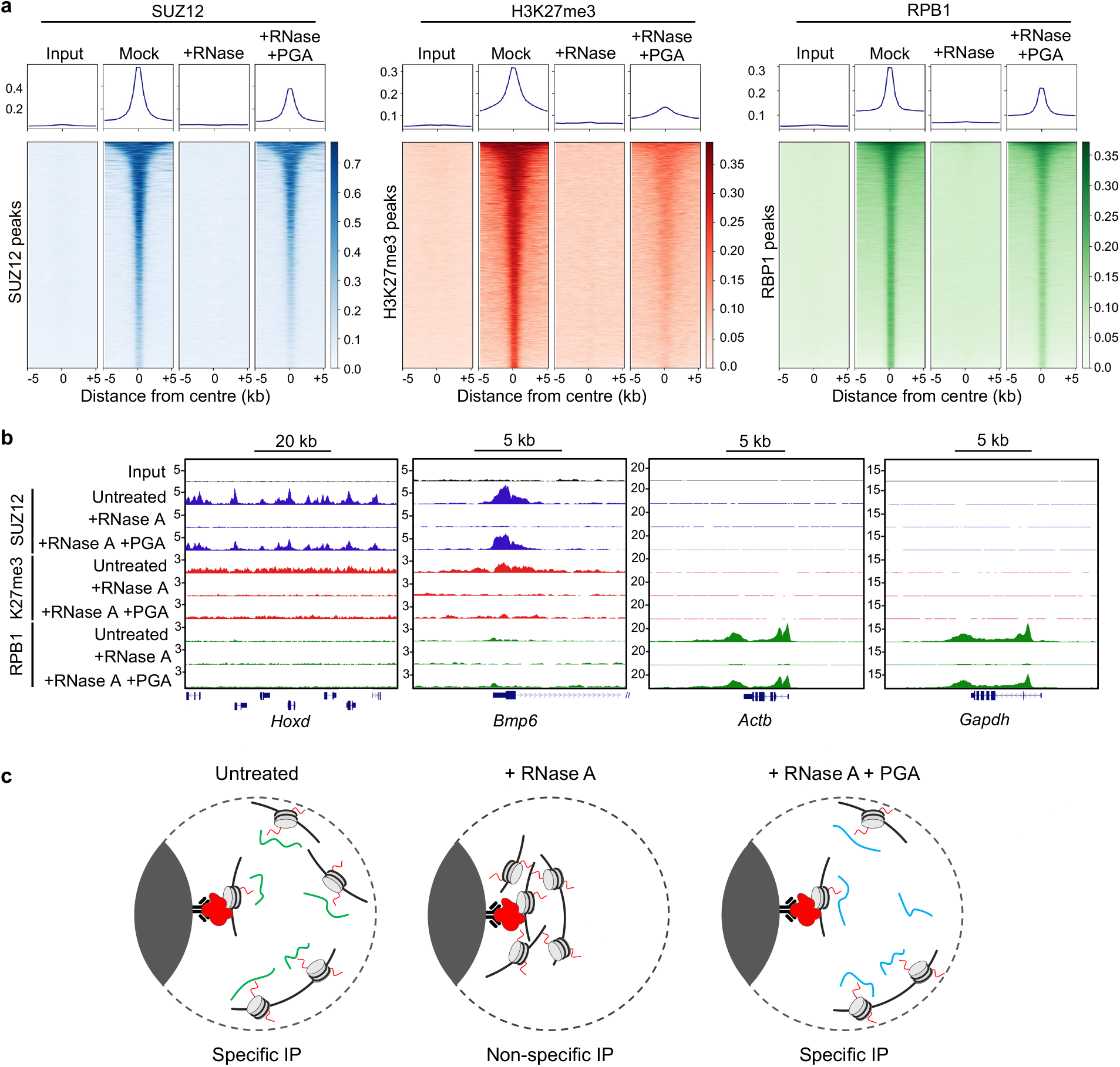
Apparent RNA-dependent PRC2 occupancy in rChIP experiments is due to non-specific chromatin IP. **a**. Metagene plots (above) and heatmaps (below) of normalized reads (per million total reads) for input and SUZ12, H3K27me3 and RBP1 rChIP samples from mock, RNase A, or RNase A and PGA treated sonicated crosslinked mESC extracts. For each epitope, normalised reads are shown centred on peaks called in any one of the three conditions. In each heatmap, normalised reads are indicated by colour, according to the scales on the right. Peaks are ordered by the average number of normalised reads. **b**. Genome browser snapshots of the *Hoxd* cluster, *Bmp6, Actb* and *Gapdh* for the samples shown in a. The y-axis shows normalised reads (per million total reads). **c**. Model: Left: In the presence of RNA, sonicated chromatin fragments are held in solution allowing for specific IP of PRC2-bound fragments. Centre: In samples treated with RNase A, non-specific fragments co-precipitate, reducing the relative enrichment of PRC2-bound genomic regions. This effect resembles the theoretical depletion of RNA-dependent PRC2 chromatin binding events. Right: The anionic polymer PGA maintains chromatin solubility in RNase A treated samples, thereby restoring IP specificity and the detection of PRC2-bound genomic regions.

In our hands, RNase A also strongly reduced the ChIP-seq signal for RBP1, and this too was substantially restored upon addition of PGA (Figs. 4a and 4b). However, although much reduced, RBP1 occupancy was still evident at active genes in RNase A-treated samples (Fig. 4b), especially in samples sonicated with the extended number of cycles used by Long et al (Fig. S1).

Taken together with our other data, we conclude that the apparent loss of PRC2 chromatin occupancy caused by RNase A in rChIP experiments is due to the non-specific precipitation of chromatin that masks the enrichment of occupied genomic regions (Fig. 4c). In the presence of RNA, sonicated chromatin fragments are held in solution allowing for specific IP of fragments bound by PRC2 or by other proteins of interest. In samples treated with RNase A, chromatin precipitates out of solution and thus non-specific fragments co-precipitate with the fragments bound by the protein of interest, reducing their relative enrichment. The anionic polymer PGA maintains chromatin solubility in RNase A-treated samples, thus restoring IP specificity and the detection of bound genomic regions.

## DISCUSSION

We report that experiments performed using rChIP methodology do not support a model in which RNA bridges PRC2 and chromatin. Rather than reducing the PRC2 ChIP-seq signal by severing an RNA bridge connecting PRC2 to chromatin, RNase A treatment of sonicated cell extracts causes non-specific chromatin precipitation, thereby diminishing the signal by increasing the background. This simple explanation also accounts for the loss of the H3K27me3 ChIP-seq signal upon RNase A treatment, and for the restoration of both the PRC2 and H3K27me3 ChIP-seq signals by the addition of the anionic polymer PGA, neither of which can be explained by an RNA bridging model. Our results agree with those of a parallel independent study by Healy and colleagues ^30^ who also demonstrate loss of ChIP specificity upon RNase A treatment when following the protocol of Long et al.

Our results are consistent with previous observations that RNase A treatment precipitates chromatin from sonicated cell extracts and that this could be rescued by the addition of PGA or other negatively charged polymers, or by deleting positively charged histone tails ^28,29^. Taken together with these previous studies, our results suggest that RNA prevents electrostatic aggregation of chromatin fragments in sonicated cell extracts. This interpretation is also consistent with the suggestion of Healy et al. ^30^ that the absence of salt in the lysis buffer used by Long et al. contributed to the apparent sensitivity of chromatin occupancy to RNA degradation.

Neither we nor Healy et al. have been able to reproduce the finding of Long et al. that RNase A has no effect on the ChIP-seq signal for RBP1, although our immunoblotting, ChIP-qPCR and ChIP-seq data indicate that the detrimental effect of RNase A treatment on IP specificity is less severe for RBP1 than for PRC2 or H3K27me3. We postulate that the milder effect on RBP1 is because active gene promoters contain nucleosome free regions ^31–33^ and thus have a lower density of charged histones.

Our demonstration that the apparent loss of PRC2 chromatin occupancy caused by RNase A treatment of cell extracts is an artefact of non-specific chromatin precipitation and that RNA degradation has little effect on PRC2 in conditions that maintain chromatin solubility undermines support for a model in which RNA bridges PRC2 and chromatin in cells. Although there is no evidence for such RNA bridges, we cannot rule out that RNA species protected from RNase A-mediated degradation perform this function. However, it is not necessary to invoke an RNA bridge model to explain PRC2 chromatin occupancy, which is dependent on recognition of H2AK119ub by JARID2 and of GC-rich DNA by PCL1-3 ^10,11^.

We also cannot rule out other mechanisms through which RNA may promote PRC2 chromatin association or activity. Long et al. reported that treatment of cells with the RNA polymerase II inhibitor triptolide depletes PRC2 from its target genes, but this effect was not observed in similar experiments by others ^34^ and experiments performed by Long et al. may have been impacted by the chromatin precipitation that occurs in RNA-depleted lysates prepared in hypotonic buffer ^28,30^.

We conclude that the apparent loss of PRC2 chromatin occupancy observed upon RNase A treatment of cell extracts in rChIP experiments does not reflect severing of an RNA bridge but is instead a symptom of non-specific chromatin precipitation. This finding highlights that ChIP-seq experiments measure enrichment of specific sequences over the genomic background and that an apparent loss of signal can instead reflect an increase in the background. We note that although the rChIP methodology produces experimental artefacts in its current form, it could still provide a means to identify RNA-dependent chromatin occupancy if performed in conditions that prevent chromatin precipitation upon RNA degradation.

## METHODS

### Cell culture

E14 mESC (gift from Helen Rowe) were cultured on 0.1% gelatin coated dishes in KO-DMEM (ThermoFisher, 10829018), 10% FBS (ThermoFisher, A3160401), non-essential amino acids (ThermoFisher, 11140035), 2 mM L-glutamine (ThermoFisher, 25030–024), 50 μM 2-mercaptoethanol (ThermoFisher, 31350010), 100 U/mL penicillin-streptomycin (ThermoFisher, 15140–122), 1 mM sodium pyruvate (ThermoFisher, 11360039) and 1000 U/mL leukemia inhibitory factor (Stemgent, 03-0011-100). The cells tested negative for mycoplasma (Lonza, LT07-701).

### rChIP

rChIP methodology was performed as described ^27^. E14 cells were cultured on 15 cm plates until 80% confluency. Cells were harvested and washed in PBS before crosslinking in 1% formaldehyde in PBS for 10 mins at room temperature (RT). Crosslinking was quenched by addition of 1/20^th^ volume 2.5 M glycine followed by rotation for 10 mins at RT. Cells were then washed twice in ice cold PBS, flash frozen and stored at −80°C in 1×10^7^ cell aliquots. Cells were resuspended in 500 μl lysis buffer (50 mM Tris-Cl pH 8.1, 10 mM EDTA, 0.5% SDS, plus protease inhibitor (cOmplete, EDTA-free Protease Inhibitor Cocktail (Roche, 04693132001) per 1.5 ×10^7^ cells. Cells were lysed on ice for 10 minutes prior to sonication for either 20 cycles (replicate 1) or 40 cycles (replicate 2) of 30s on / 30s off with a Bioruptor UCD-200 (Diagenode) at maximum power. Lysates were centrifuged at 17,000g for 10 minutes at 4°C, the supernatant collected and pre-cleared with Pierce protein A/G magnetic beads (ThermoFisher 88803) in IP buffer (16.7 mM Tris-HCl pH 8.1, 1.2 mM EDTA, 167 mM NaCl, 1% Triton X-100, cOmplete protease inhibitor). Cell extract equivalent to 3 ×10^6^ cells were diluted 1:5 in IP buffer with 2% taken and stored at −20°C as input. 2.5 μl anti-H3 (2.5 μg; Abcam, 1791), 2.5 μl anti-H3K27me3 (2.5 μg; Abcam, ab192985), 2.5 μl anti-SUZ12 (Cell Signaling, 3737) or 2.5 μl anti-RPB1 (Cell Signaling, 14958) antibody was added to the extracts, followed by incubation at 4°C overnight. For RNase A treated conditions, 5 μg/ml RNase A (Thermo Scientific, EN0531) was added together with the antibody, with or without the addition of 400 μg/ml PGA (Sigma-Aldrich, P4761). Following overnight incubation, 25 μl of Pierce protein A/G beads were added and incubated for 1 hr at RT. Beads were captured with a magnetic rack and washed twice with 1 ml of each of the following buffers: (20 mM Tris-Cl pH 8.0, 2 mM EDTA, 150 mM NaCl, 0.1% SDS, 1% Triton X-100), high salt buffer (20 mM Tris-Cl pH 8.0, 2 mM EDTA, 500 mM NaCl, 0.1% SDS, 1% Triton X-100), LiCl buffer (10 mM Tris-Cl pH 8.0, 1 mM EDTA, 250 mM LiCl, 1% sodium deoxycholate, 1% IGEPAL CA-630 (Sigma-Aldrich, 6741), TE (10 mM Tris pH 8.0, 1 mM EDTA). Chromatin was eluted in 120 μl elution buffer (100 mM sodium bicarbonate, 1% SDS) for 20 minutes at RT. Input samples were thawed, topped up to 120 μl with elution buffer and incubated in parallel. Crosslinks were reversed through addition of 200 mM NaCl and incubation at 65°C overnight. DNA was purified through phenol-chloroform-isoamyl extraction and ethanol precipitation with addition of GlycoBlue (ThermoFisher, AM9516). DNA was resuspended in 30 μl 10 mM Tris-HCl, quantified by Qubit and the size range determined on a DNA Bioanalyser (Agilent). Specific DNA sequences were quantified by qPCR. Libraries were prepared from 2.5 ng of each sample using either the KAPA HyperPlus Kit (replicate 1: Roche/KAPA Biosystems, KK8512) or NEBNext Ultra II DNA Library Prep Kit for Illumina (replicate 2: NEB, E7645S), according to the manufacturers’ protocols. DNA libraries were pooled and sequenced with an Illumina NovaSeq (150 bp, paired end) by Novogene UK.

### rChIP data analysis

Sequencing analysis was performed using the NF-core ChIP-seq pipeline ^35^. Briefly, rChIP-seq reads were trimmed using trimgalore! and aligned to the reference genome GRCm38 using BWA (V 0.6.0) ^36^. Duplicates were removed using picard-tools, blacklisted regions were filtered and removed using SAMtools (1.18) ^37^ and peaks called with MACS2 (3.0.0b2) ^38^ with the --broad option. Normalised bigwigs scaled to 1 million mapped reads were generated using BEDTools (2) ^39^ and visualised using the UCSC genome browser. Metaplots and heatmaps showing occupancy across all the set of peaks called for each factor were generated with the computeMatrix and plotProfile/plotHeatmap functions from deepTools ^40^.

### Separation of extracts into soluble and insoluble fractions

Extracts were prepared as above except lysates were sonicated for 12 cycles of 30s on / 30s off, using a Bioruptor Pico (Diagenode). After incubation for 1 hr at 4°C and then 1 hr at RT with or without RNase A or PGA, extracts were centrifuged at 17,000g for 10 mins at 4°C to separate soluble (supernatant) from insoluble (pellet) material.

### Co-immunoprecipitation

Extracts were prepared as for rChIP except that lysates were sonicated for 12 cycles of 30s on / 30s off, using a Bioruptor Pico (Diagenode). Sonicated extracts were diluted 1:5 in IP buffer (16.7 mM Tris-HCl pH 8.1, 1.2 mM EDTA, 167 mM NaCl, 1% Triton X-100, cOmplete protease inhibitor) with 2% extract (relative to the volume of an individual IP) removed as input. Extracts were split equally and incubated with 2.5 μl anti-H3 (2.5 μg; Abcam, 1791) 2.5 μl anti-SUZ12 (Cell Signaling, 3737) and 2.5 μl anti-RPB1 (Cell Signaling, 14958) antibody overnight at 4°C with or without RNase A before the addition of Pierce protein A/G beads and incubation for 1 hr at RT. Beads were washed according to rChIP methodology and resuspended alongside inputs in NuPAGE LDS Sample Buffer (4X) (Invitrogen, NP0007) and incubated at 70°C for 10 minutes.

### SDS-PAGE

Extracts were centrifuged at 16,000g for 10 mins at 4°C to separate soluble (supernatant) from insoluble (pellet) material. The pellet was resuspended in an equal volume of 5:1 IP:lysis buffer. LDS sample buffer was added and proteins were resolved on 4-12% NuPAGE Bis-Tris precast gels (Life Technologies, NP0322) alongside PageRuler Plus Prestained Protein Ladder (Thermo Scientific, 26619). Proteins were visualised using Oriole fluorescent gel stain (Bio-Rad, 161-0496) and a BioRad ChemiDoc MP.

### Immunoblotting

Proteins were transferred onto nitrocellulose membranes (Amersham, 10600002) in Tris-Glycine transfer buffer with 20% methanol. Membranes were blocked in TBS+0.1% Tween-20 (TBS-T) + 5% milk and then incubated overnight at 4°C with the following primary antibodies in TBS+5% milk: anti-RPB1 (1:1000; Cell Signaling, 14958), anti-EZH2 (1:1000; Cell Signaling, 5246), anti-SUZ12 (1:1000; Santa Cruz, sc-271325), anti-Beta-Tubulin (1:1000; Abcam, ab6046), anti-β-Actin (1:1000; Cell Signaling 4967S), anti-H3K9me3 (1:1000; Abcam, ab8898) anti-H3K27me3 (1:1000; Abcam, ab192985), anti-H3K27ac (1:1000; Abcam, ab4729) and anti-H3 (1:2000; Abcam, 1791). Membranes were washed 3x in TBS-T, incubated with HRP-conjugated goat anti-rabbit (Dako, P0448) or rabbit anti-mouse (Dako, PO260) secondary antibodies, washed again in TBS-T and proteins visualised with ECL (Biorad, 170-5061) and an Image Quant 800 (Amersham).

### DNA and RNA purification and agarose gel electrophoresis

DNA was extracted from supernatants by addition of phenol:chloroform:isoamyl alcohol (25:24:1). The aqueous phase was isolated through centrifugation in MaXtract high density tubes (Qiagen, 50-727-738). DNA was precipitated at −20°C by the addition of 100% ethanol, NH_4_OAc and GlycoBlue coprecipitant, washed in 80% ethanol and resuspended in TE. Equal volumes of DNA were loaded onto a 1.5% agarose gel and visualised using SYBR Safe nucleic acid gel stain (Invitrogen, S33102) and the UVP BioDoc-It Imaging System. RNA was isolated from sonicated lysates using TRIzol LS Reagent (ThermoFisher, 10296028) according to the manufacturer’s instructions. Equal volumes of RNA were resolved on Novex TBE-urea gels (ThermoFisher, EC6865) and visualised as for DNA.

### qPCR

qPCR was performed with SYBR Green Master Mix (Bio-Rad, 1725272) and the primers listed below, in a QuantStudio 5 Real-Time PCR System (ThermoFisher) using the default program. The amount of each DNA sequence in ChIP samples were calculated relative to input using the 1′Ct method and mean and standard error calculated from triplicate experiments. The significance of differences in % input and ratio to *Gapdh* were estimated using unpaired t-tests.

### Primers (5’-3’)

*Hoxd11* F – GGCCGAGGGTTCTCCCCCTT

*Hoxd11* R – CCTCCCTCCCCCACCACCAG

*Actb* F – AGGAGCTGCAAAGAAGCTGT

*Actb* R – CCGCTGTGGCGTCCTATAAA

*Bmp6* F – AGCCGCCTCTGAGGGTTC

*Bmp6* R - GCCAGGTGTGTCCTAGGCAG

*Gapdh* F – CCCACTCCGCGATTTTCA

*Gapdh* R – CTCTGCTCCTCCCTGTTCCA

## Data availability

ChIP-seq data have been deposited at Gene Expression Omnibus (GEO) and are available via accession number GSE240380.

## Acknowledgements

We thank Chen Davidovich, Tom Cech and John Rinn for discussions and for sharing unpublished data. Thanks to Yicheng Long and John Rinn for sharing the full rChIP protocol. This work was funded by a Cancer Research UK (CRUK) UCL Centre (C416/A18088) PhD studentship and a Biotechnology and Biological Sciences Research Council grant (BB/W008750/1).

## Contributions

A.H.H: Investigation, Formal analysis, Data curation, Writing – original Draft, Writing – Review and Editing. R.G.J: Conceptualisation, Methodology, Writing – original Draft, Writing – Review and Editing, Supervision.

## Ethics declarations

Competing interests: The authors declare no competing interests.

**Figure S1.**
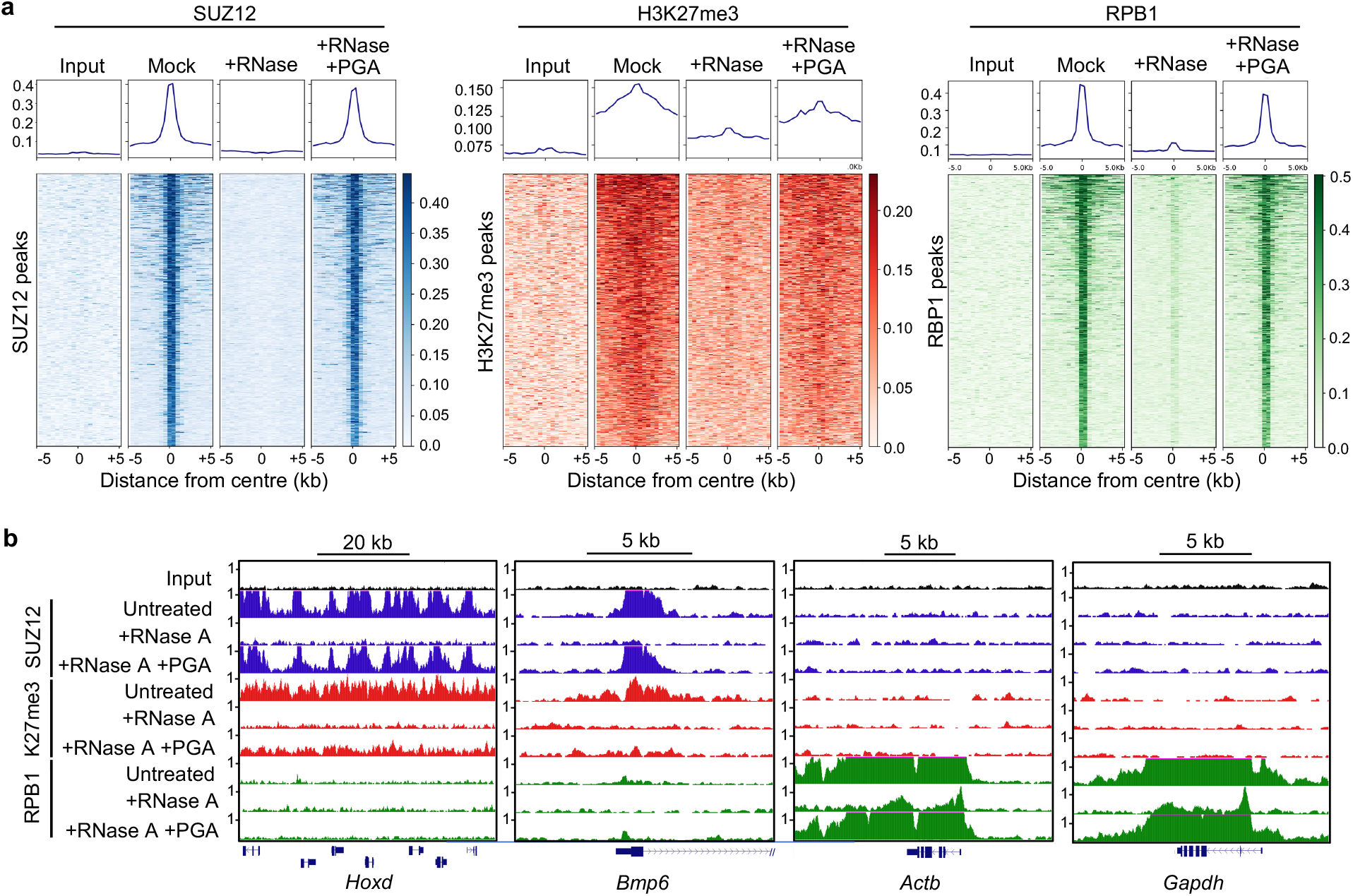
Detection of RBP1 chromatin occupancy by rChIP is less affected by RNase A treatment than PRC2 chromatin occupancy. **a**. As Figure 4a, except for a second independent replicate performed with a greater number of sonication cycles (40 vs 20) that more closely matches that performed by Long et al. **b**. As Figure 4b, but with lower maximum y-axis values to visualise RBP1 occupancy more clearly at *Gapdh* and *Actb* in RNase A treated samples.

